# Social or non-social? An exploratory approach to study inequity aversion in primates

**DOI:** 10.64898/2026.06.10.731392

**Authors:** Aurore Serda, Hélène Meunier, Charlotte Canteloup

## Abstract

Inequity aversion, the sensitivity to inequitable outcomes or processes, has been widely studied in nonhuman primates since 2003. Yet, findings remain debated as some could be explained by alternative mechanisms such as frustration or loss aversion, so-called ‘individual contrasts’. Previous work on nonhuman animals has restricted the definition of inequity aversion to the social domain. Here, we propose distinguishing between two different forms: a socially driven form, which depends on comparison with another individual, and an individually based form, caused by a discrepancy between one’s own effort and the expected outcome. Using a task-based methodology, we manipulated both effort and the amount of reward to test for the existence of those different forms and distinguish them from individual contrasts. We presented seven Tonkean macaques (*Macaca tonkeana*) and three brown capuchins (*Sapajus apella*) with low and high effort tasks associated with low and high reward quantities. To test for the presence of individually based inequity aversion, subjects were tested alone in an individual phase. In some of the trials, they were rewarded less than they deserved for the task. To test for socially based inequity aversion, two individuals performed the same task in a social phase but, in some trials, one received a higher-value reward than the other. We recorded the latency to engage in the task as an indicator of reluctance. In the individual phase, macaques were slower to engage with inequitably rewarded tasks, a pattern consistent with an individually based expectation of equity. By contrast, capuchins were faster in this context, suggesting that their responses were more likely driven by individual contrast effects. In a context of social inequity, both species slowed engagement in tasks with inequitable rewards, suggesting an aversion to socially based inequity. These results demonstrate that such a methodology provides a means to study non-social components of inequity aversion. Future studies, conducted on larger and more diverse samples, with rigorous motivational controls and explicit tests looking at the understanding of the link between task and reward, are needed to confirm whether nonhuman primates can display inequity aversion independently of social comparison.

## Introduction

Many cooperative behaviours such as hunting (Boesch, 1994), defending (Port, Kappeler & Johnstone, 2011) and parenting (Burkart & van Schaik, 2010) have been considered more advantageous for some social primate species when shared among group members (Kappeler & van Schaik, 2006; Silk, 2007). Most research focuses on how and by whom those actions are shared, but fewer studies investigate the rules underlying the distribution of the resulting rewards. For example, some observational data from hunting events suggest that chimpanzees and capuchins who actively contribute to the hunt get a bigger share (Boesch, 1994, 2002; Rose, 1997). A remaining open question is: do members of the hunt take into account each individual’s involvement when reacting to the reward they get? In other words, do they consider equity, i.e. one’s ratio of outcomes to input being equal to that of conspecifics (Adams, 1965; Vale & Brosnan, 2022), when responding to the situation?

The ‘cooperation hypothesis’ (Brosnan, 2011) suggests that the aversion to violations of equity-known as *inequity aversion*- may be a key mechanism promoting long-term success in cooperation. In particular, this capacity may have helped stabilize cooperative exchanges by enabling individuals to choose optimal partners. By most of its definitions, *inequity aversion* is a social mechanism. The Encyclopedia of Animal Cognition and Behavior states that ‘inequity aversion in animals is a response to outcomes that are less than, or more than, what a social partner receives’ (Vale & Brosnan, 2022). Robustly documented in humans (e.g. Engel, 2011; Henrich et al., 2004; Tabibnia & Lieberman, 2007), the existence of *inequity aversion* remains debated in nonhuman animals (Bräuer & Hanus, 2012; Oberliessen & Kalenscher, 2019; Ritov et al., 2024). Numerous studies have reported negative reactions to differences in reward distribution across a variety of nonhuman primates, hereafter ‘primates’ (e.g. Brosnan & de Waal, 2003; Talbot et al., 2018; van Wolkenten et al., 2007; Verspeek & Stevens, 2023; Yasue et al., 2018). Different methodologies have been used to question the underlying mechanisms of these reactions (e.g. Brosnan, Freeman & De Waal, 2006; Jensen, Call & Tomasello, 2007; Fletcher, 2008; van Leeuwen, Zimmermann & Ross, 2011; Campbell et al., 2020), the pioneering paradigm being the token exchange task (Brosnan & de Waal, 2003). In this protocol, two individuals are trained to return a token to an experimenter in exchange for a food reward. Under equitable conditions, both monkeys receive the same reward for the exchange. But under inequitable conditions, one of the monkeys receives a higher-value reward for the same effort. The authors showed that brown capuchin monkeys (*Sapajus apella*) often refused the exchange when their partner received a more attractive reward for the same effort (Brosnan & de Waal, 2003). They interpreted this behaviour as a possible indicator of *inequity aversion*. Since then, research on the subject has expanded, and other experiments have challenged this conclusion, proposing alternative explanations for the results obtained (e.g. Fontenot et al., 2007; Bräuer, Call & Tomasello, 2009; Freeman et al., 2013; Schweinfurth & Call, 2021).

One alternative mechanism that is often mentioned is that of *individual contrasts* (Ind-C), which, most often, refer to frustration and loss aversion (Wynne, 2004; Fontenot et al., 2007; Silberberg et al., 2009; Oberliessen & Kalenscher, 2019; Ritov et al., 2024). For example, subjects might react negatively during the token exchange task simply because a better reward was not available to them. This negative reaction might not depend on the fact that this inaccessible reward was specifically given to the partner. If subjects react in the same way when the better reward is simply placed inside an adjacent empty cage, this would indicate that frustration is a more parsimonious explanation than *inequity aversion*. Several studies have highlighted such effects, thereby calling into question the conclusions of earlier research (Fontenot et al., 2007; Kaiser et al., 2012; McAuliffe et al., 2015).

Moreover, most researchers consider the social dimension of *inequity aversion* to be an integral part of its definition. This has led them to interpret all reactions under individual control conditions as indicative of alternative mechanisms such as Ind-C. Conversely, in human psychology, *inequity aversion* is not necessarily considered to be a purely social mechanism, but one that could actually be triggered by an internal comparison alone (Kelley, 1959; Pritchard, 1969; McAuliffe et al., 2013). This idea of an internal point of reference has been present in the field of human psychology for decades, both with Kelley’s (1959) ‘Comparison Level’ or with Pritchard’s (1969) ‘internal standard’. The latter has been defined as ‘the amount of outcome Person perceives as being commensurate with his own inputs, without regard to any comparison person’ (Pritchard, 1969, p.205). Studies show that both children (McAuliffe et al., 2013) and adults (Pritchard, Dunnette & Gorgenson, 1972) show signs of *inequity aversion* even under non-social conditions.

We propose that, in the case of primates as well, reactions to inequity may result from two partially distinct processes: (i) a Socially based Inequity Aversion (Soc-IA) which is based on comparison with a conspecific, and (ii) an Individually based Inequity Aversion (Ind-IA) triggered by a mismatch between an individual’s own input and the outcome obtained. What distinguishes Ind-IA from Ind-C, i.e. frustration or loss aversion effects, is the concept of ‘merit’, whereby the outcome obtained should correspond to the effort expanded. For instance, a macaque may experience frustration when denied access to an infant, regardless of any social comparison or personal actions. This would be consistent with an individual contrast (Ind-C) effect. However, if the individual has previously invested substantial affiliative effort, such as prolonged grooming of the mother, the mismatch between effort and outcome may instead reflect an Individually based Inequity Aversion (Ind-IA), regardless of whether others are rewarded differently. In contrast, a socially driven inequity response (Soc-IA) would occur if another individual were granted access to the infant even though their level of investment was similar or lower. In primates, Ind-IA is never questioned and remains conflated with Ind-C. Previous studies have not examined whether reactions consistent with Ind-C were task or effort specific (Oberliessen & Kalenscher, 2019; Ritov et al., 2024), even though these two concepts may not actually be interchangeable (Serda, Canteloup & Meunier, preprint).

Distinguishing between these mechanisms might be essential. A process distinct from both Ind-C and Soc-IA might indeed offer strong evolutionary advantages. It could allow the integration of more information than Ind-C, while still being less cognitively demanding than Soc-IA. For example, evaluating socially mediated equity in the sharing of the loot of a group hunt, would require a heavy cognitive load for individuals. They should indeed be able to track each conspecific’s actions, evaluate the quantity of each partner’s share, and compare those according to their identities, while monitoring most of the hunt and the final distribution of meat. A simpler way to avoid being slighted is to rely on an internal standard of what one’s own actions in a hunt should generally yield, based on one’s identity, efforts, and needs. Established by previous experiences, such a standard would be easily accessible and tailored to each individual’s situation in their social group. It would also go beyond general Ind-C, which would not account for individual participation or promote cooperative group dynamics in the same way.

A major challenge in testing for Ind-IA is that it requires establishing a clear relationship between effort and expected reward. To date, research conducted on inequity has generally not taken into account the value of the outcome in relation to the individual input for the subjects. Indeed, when focusing solely on Soc-IA, the methodology emphasizes the violation of the compared outcome between subjects, not within a single subject. In the same way, controlling for Ind-C alone does not imply a methodology establishing the value of the input. Some researchers have examined the scale of this effort and demonstrated its importance in highlighting inequity averse responses (van Wolkenten, Brosnan & de Waal, 2007; Massen et al., 2012; McAuliffe, Shelton & Stone, 2014). But even in those studies, effort was not examined in direct comparison with the reward it is supposed to help achieve.

The present study aims to address this gap by proposing a new paradigm. Our goal is to establish a proof of concept that such a design makes it possible to identify a specific mechanism – analogous to humans’ *inequity aversion* based on internal standards (Ind-IA). This should also allow comparison with socially based *inequity aversion* (Soc-IA) and distinguish *individual contrast* effects (Ind-C). To this end, this methodology relies on various tasks that had been taught to the monkeys, and that always yielded different results: completing the task that required significant effort (‘high-effort task’) always resulted in a substantial reward, whereas completing the task that required little effort (‘low-effort task’) always resulted in a smaller reward. This allowed us to either give subjects the reward they deserved (equity) or to reward them less than they deserved (inequity). In an individual context, only the internal standard of reward regarding the effort was available, whereas in a social context, a partner’s relative reward served as a point of comparison. The novelty of this approach lies in the fact that it involves two tasks with two distinct, equitable but not equal, rewards.

We conducted this study on capuchin monkeys, a species that has already been tested in several studies of *inequity aversion* in primates (Brosnan & de Waal, 2003; Schweinfurth & Call, 2021) as well as on Tonkean macaques (*Macaca tonkeana*), which had not yet been tested in this context. This provided a baseline for examining the species-specific aversion to inequity as well as for studying a species that had never been involved in such research. We predict that if subjects are sensitive to Ind-IA, they will be reluctant to engage in tasks rewarded inequitably but not in those rewarded unequally, even in the absence of a partner. They would do so by reacting negatively to a low reward for a high-effort task, but not (or less) to a low reward for a low-effort task. We further expect that these responses may be modulated by the social context, leading to stronger reactions in the presence of inequity relative to a conspecific. If subjects do indeed respond to Ind-IA, this would support the proposed paradigm and challenge the current tendency to dismiss certain forms of Ind-C as alternative explanations. Conversely, if responses are solely driven by Ind-C or Soc-IA, this could highlight the importance of social comparison or invalidate our experimental protocol.

## Materials & Methods

### Ethical statement

All procedures were non-invasive. The animals’ participation was always voluntary, subjects were never restrained or forced to interact with the tasks. No food or water restriction was used. This research was approved by the ethical committee of the Primate Center of the University of Strasbourg (SBEA), which is authorized to house non-human primates (registration n° B6732636). The research further complied with the EU Directive 2010/63/EU for animal experiments. Generative AI was used for coding assistance only.

#### 1. Subjects and housing

The study took place between January and March 2025. It involved ten adult male subjects, including seven Tonkean macaques and three brown capuchins, aged between 7 and 17 years (Table 1). All subjects were housed in only male social groups ranging from two to six individuals at the Simian Laboratory Europe - University of Strasbourg.

**Table 1.** : **Subject identity, age, species, group size, and housing conditions.**

| Subject | Age (years) | Species | Group Size | Housing |
| --- | --- | --- | --- | --- |
| Walt | 17 | *Macaca tonkeana* | 6 | Large naturalistic outdoor enclosure, ~1364m<sup>2</sup> outdoor + indoor access |
| Yang | 15 |  |  |  |
| Olli | 14 |  |  |  |
| Barnabé | 7 |  | 3 | Small concrete outdoor enclosure, ~32m<sup>2</sup> outdoor + indoor access |
| Abricot | 10 |  |  |  |
| César | 12 |  | 2 | Small concrete outdoor enclosure, ~32m<sup>2</sup> outdoor + indoor access |
| Alaryc | 11 |  |  |  |
| Franklin | 9 | *Sapajus apella* | 3 | Small concrete outdoor enclosure, ~8m<sup>2</sup> outdoor + indoor access |
| Kiri | 14 |  |  |  |
| Brugnon | 12 |  |  |  |

Housing conditions varied across groups. Tonkean macaques were either housed in large naturalistic wooded outdoor enclosures, or in smaller concrete-floored outdoor enclosures with basic enrichment (platforms, climbing structures). Brown capuchins were housed exclusively in small outdoor concrete enclosures equipped with wooden structures and some enrichment objects (e.g., toys, ropes). All groups had continuous access to indoor facilities (Table 1). Water was available *ad libitum*, and animals were fed daily with pellets. Fresh fruits and vegetables were provided once a week.

Eight individuals (Abricot, Alaryc, César, Franklin, Kiri, Olli, Walt, Yang) participated in the ‘individual phase’ of the experiment. Among these, four (Alaryc, César, Franklin, and Kiri) also served as subjects in the ‘social phase’, where they were paired with a conspecific partner. Four animals (Abricot, Barnabé, Kiri and Brugnon) served as model partners but three of them were not tested themselves in the ‘social phase’ (Abricot, Barnabé, and Brugnon). All individuals were tested either directly in their home environment or in a separate dedicated space that was part of their home environment.

A summary of subjects, species, age, group size, and housing conditions is presented in Table 1.

#### 2. Tasks and apparatus

Two tasks were used to provide different effort requirements:

### A low-effort task

A simple target consisting of a 4cm diameter wooden ball, attached to the top of a metal rod, which the subject had to touch with their hand through the mesh. Completion of this task provided one reward (one raisin).

### A high-effort task

A custom-designed puzzle (32x5x27cm, Figure 1) requiring subjects to push a wooden ball (identical to the one used for the low-effort task) with their hand through a vertical series of four compartments to obtain rewards (four raisins). The puzzle was made of opaque PVC and featured a transparent acrylic panel on the experimenter’s side to allow for visual monitoring. Metal rods were present on the side facing the subject to prevent direct grasping of the ball, thereby ensuring that the task had to be performed. The subject had to push the wooden ball through each level until it reached the final level and fell into a transparent receptacle. Once the ball was retrieved by the experimenter, they delivered the rewards.

**Figure 1.**
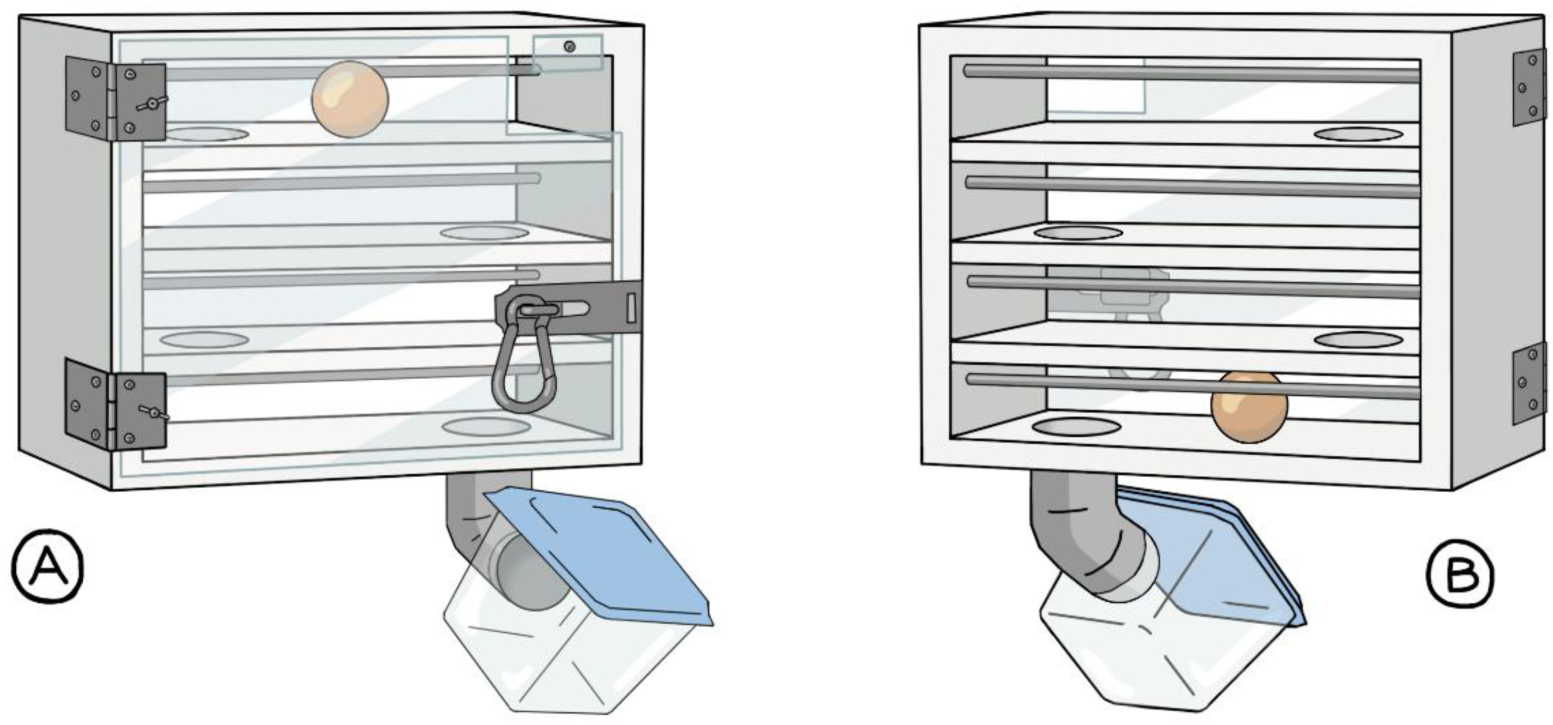
: view of the apparatus used as the high-effort task. (A) From the experimenters’ side. (B) From the subject’s side. Illustration by Manon Serda.

#### 3. Training

All subjects underwent a training phase to ensure that they were able to reliably perform the low-effort and high-effort tasks prior to the tests. Subjects validated this phase when they were able to complete the entire sequence of tasks during twelve consecutive trials, in two consecutive sessions, without any assistance. Then, they underwent several training sessions aimed at associating each task with each reward. Subjects performed 180 trials of the high-effort task over fifteen sessions (with the exception of two of our subjects who performed 168 trials spread over fourteen sessions and one subject who performed 144 trials spread over twelve sessions) to ensure that they reliably expected to receive four raisins for this task. Since our subjects were accustomed to receiving one single reward in response to a target command, we conducted only 48 trials spread over four sessions to associate this specific target with a single raisin.

To control for potential confounding factors related to satiety and reward evaluation, two additional pretests were conducted. First, a satiety test was used to verify that each subject was willing and able to consume the maximum number of raisins corresponding to the highest possible reward during a single session. Second, a quantity discrimination test ensured that all individuals preferred the greater amount of reward and were able to reliably distinguish between a reward of one *versu*s four and four *versus* eight, by consistently pointing to the larger amount.

Only subjects meeting all criteria were included in the subsequent experimental phases.

#### 4. Procedure

At the start of each trial, the experimenter first placed the number of raisins corresponding to the test condition in the subject’s cup, in front of them but out of their reach. They always made sure beforehand that the individual was attentive to the task (i.e. looking toward the experimenter and their cup). Once the reward was clearly visible, so that the subject could anticipate the outcome associated with completing the task, the experimenter presented the task to the subject. To do this, they presented the ball-target close to the mesh (low-effort task) or inserted the ball into the top level of the puzzle (high-effort task). The trial continued until the task was completed and the reward was delivered, unless the subject stopped participating for 30 consecutive seconds. In that case, the trial was considered a refusal and the next trial began. If the subject did not begin the task within 30 seconds after presentation, the trial was also considered a refusal. If the subject did not begin to take or consume the reward within 30 seconds of its presentation, the reward was considered rejected. Each trial began immediately after the previous one, leaving only a few seconds for the subject to consume the reward and for the experimenter to prepare the next task. The food rewards, raisins, were quite appetizing to all subjects.

##### 4.1 Individual phase

In the individual phase, subjects (n=8) were tested alone in the experimental area. Each session consisted of 12 trials, with six low-effort and six high-effort tasks presented in a pseudo-randomized order (no more than two consecutive trials of the same task type were allowed within a session). No more than one session per day was conducted for a total of four sessions.

Each subject completed two equity sessions (EQ-I), followed by two inequity sessions (INE-I). This order was established to prevent the inequity trials from influencing the behaviours observed in the equity condition. During the equity sessions, as expected, subjects received one raisin for completing the low-effort task and four raisins for the high-effort task. During the inequity sessions, the task structure remained the same, but the reward for the high-effort task was deliberately reduced to a single raisin, thereby creating a reward/effort mismatch.

##### 4.2 Social phase

In the social phase, the subject (n=4) and a familiar conspecific (the model, n=4) were placed in adjacent compartments separated by a mesh panel (Figure 2). Both individuals had visual access to each other and their respective reward cups. Trials were conducted sequentially, with the model and subject alternating in completing the task, always beginning with the model. Only one individual performed a trial at a time.

**Figure 2:**
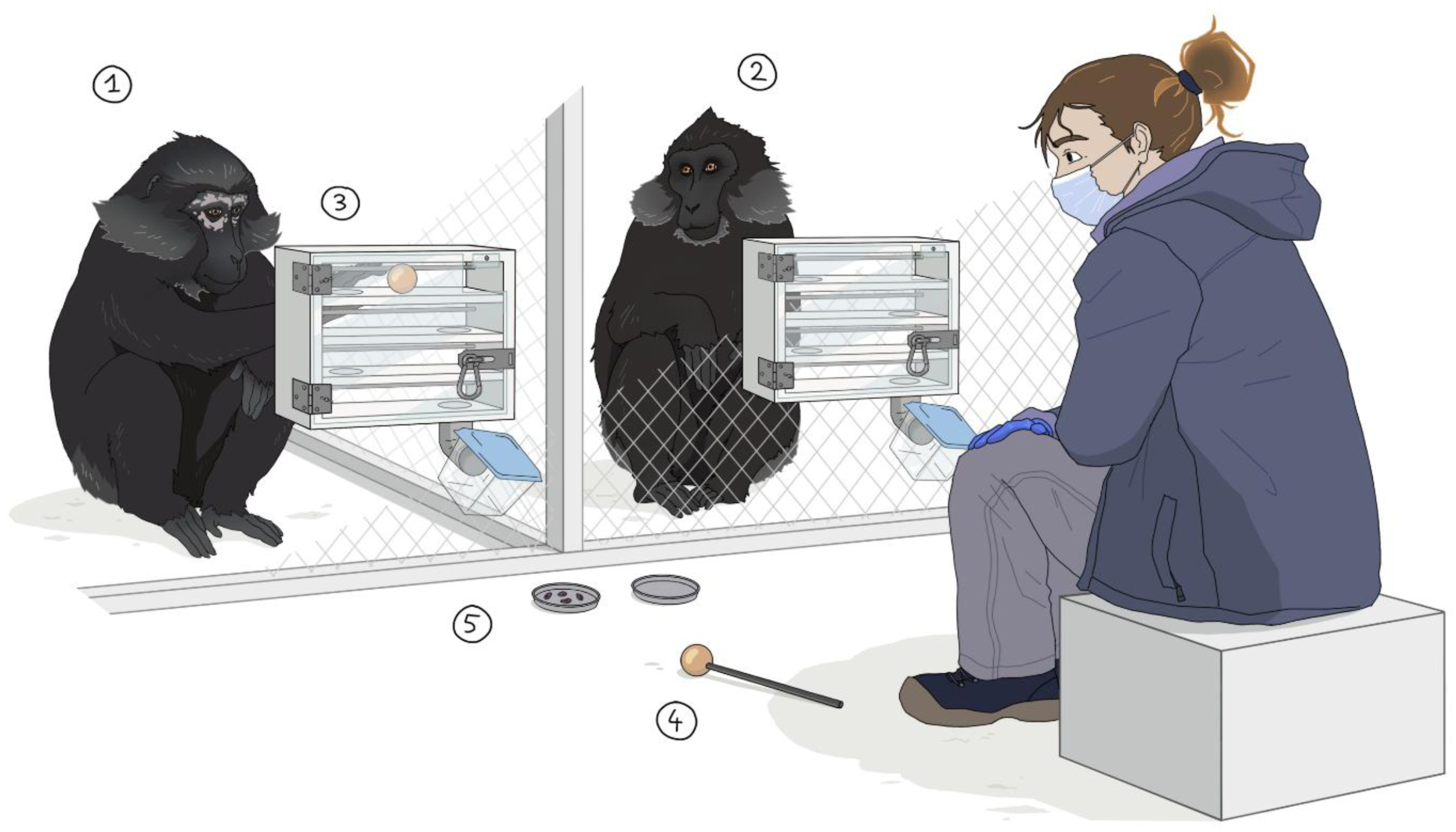
Visual representation of the social phase experimental set up. The model (1) is completing the high-effort task (3) for four raisins (5) while the subject (2) watches and waits for his turn. The target (4) on the ground will be used as the low-effort task. Illustration by Manon Serda.

As for the individual phase, each pair completed two equity (EQ-S) and two inequity (INE-S) sessions, with each session consisting of 12 trials per individual. For subjects paired with two different models, this resulted in a total of eight sessions (in no cases did any individual take on the role of subject after having been a model). The order of conditions was established as in the individual phase to prevent reward-based frustration biases from influencing the equity condition.

During the equity sessions, both the subject and the partner received a reward proportional to the task’s difficulty: one raisin for the low-effort task and four raisins for the high-effort task. In the inequity sessions, the subject continued to receive these expected rewards, while the partner always received eight raisins for completing the high effort task (and the expected one raisin for the low effort task).

The type of effort was pseudo-randomized, and the trial sequence was balanced so that the subject’s task was just as likely to follow a model trial of the same or different effort level.

#### 5. Control measures for alternative hypotheses

When designing this study, we considered several alternative hypotheses, including frustration, loss aversion, motivation, and arousal (details and results for the control of arousal can be found in Supplemental Files S1). We implemented specific controls to account for each mechanism when appropriate.

##### 5.1 Frustration

We controlled frustration at two levels:

a. <u>Frustration resulting from seeing a more valuable but out-of-reach reward</u>: we ensured that a large number of rewards was always visible to the subjects. By placing a transparent box filled with raisins in front of them throughout the sessions, we ensured that the resulting frustration was consistent across all conditions.
b. <u>Frustration resulting from observing the distribution of a higher-value reward</u>: we added two social frustration control sessions per tested pair (CTRL-F). During these sessions, the subject and the model performed the same six low-effort trials and received identical low-value rewards (one raisin), with no high-effort trials presented. We compared responses to these trials to low-effort trials of the social equity condition, where low-effort tasks alternated with high-effort tasks for which the model received a higher-value reward. This allowed us to verify whether simply witnessing a partner receiving more, even when distribution remained equitable, elicited frustration. Although this comparison focuses on the low-effort task, it indirectly sheds light on the interpretation of frustration effects more generally. A more direct control for the high-effort task could not be implemented without confounding inequity.

Such controls were not necessary during the individual phase, when no conspecific was present and no differential distribution of reward was observable.

##### 5.2 Loss Aversion

During the individual phase, we assessed loss aversion indirectly through the low-effort task. Since the sessions always included both types of tasks, receiving a low-value reward after a higher-value reward could elicit loss-averse behaviours. The inequity condition served as a control, as in this condition, the low-reward task was never alternated with a more rewarding task. Differences between these conditions could help distinguish the effects of *inequity aversion* from general loss sensitivity.

In the social phase, loss aversion was not a confounding factor, as the subject’s own reward remained constant across the equity and the inequity conditions and was never reduced.

##### 5.3 Motivation

During the social phase, the subject’s rewards remained the same across equity and inequity sessions, eliminating confounding factors related to motivation.

During the individual phase, we conducted two separate pre-test sessions before the tests began. During these sessions, subjects performed the high-effort task in exchange for a single raisin, before any association between reward and effort had been established, to serve as a baseline for motivation.

All conditions are summarized in Table 2.

**Table 2:** Experimental and control conditions. EQ = equity; INE = inequity; CTRL = control; F= frustration; R = raisin. The individual phase refers to the subject being tested alone, and the social phase refers to the partner being present.

| Acronym | Phase | Effort–reward relationship | Reward structure: (Low–High effort; Subject / Model) | Purpose |
| --- | --- | --- | --- | --- |
| EQ-I | Individual | Effort-matched | 1R-4R/ - | Baseline for individual evaluation |
| INE-I | Individual | Reward reduced for high-effort | 1R-1R/ - | Test individual *inequity aversion* |
| EQ-S | Social | Both effort-matched | 1R-4R/1R-4R | Baseline for social evaluation |
| INE-S | Social | Partner gets double reward for same effort | 1R-4R/1R-8R | Test social *inequity aversion* |
| CTRL-F | Social | Both perform low-effort for 1R | 1R/1R | Control for frustration (no reward asymmetry, no inequity) |

#### 6. Data collection and coding

All sessions were video recorded using GoPro HERO10 (GoPro Inc., San Mateo, CA, USA), and behavioural coding was performed using BORIS software (v. 8.27.10, Friard & Gamba, 2016).

For each trial, we recorded whether the subject agreed to participate (by performing the task and eating the reward) or refused (by remaining inactive for 30 consecutive seconds without interacting with the task or eating the reward). The initiation latency was defined as the time interval between the presentation of the ball and the subject’s first action, that is, the moment they touched the ball.

A subset of 20% of the video sessions was randomly selected for double coding. Inter-rater reliability was assessed using Cohen’s Kappa (all Kappa > 0.80). Coders could not be blind to experimental conditions.

#### 7. Statistical analyses

All analyses were conducted in R 4.5.0 (R Core Team, 2025) using generalized linear models (GLMs) and generalized linear mixed models (GLMMs) using package glmmTMB (Brooks et al., 2017). We explored seven different models per species, to account for seven different hypotheses, listed in the table below (Table 3). The response variable in all models was the latency to initiate the task (in seconds). Fixed effects included condition (EQ, INE and CTRL-F), phase (individual, social), preceding task (high or low effort), session (1-4) and trial numbers (1-12) according to the specific hypothesis tested. GLM(M)s were chosen because they allow for complex interactions, can handle repeated measures (controlling for within-subject correlation), and, if needed, may adapt to heterogeneous residual variance. Because of our small sample size, non-parametric alternatives were considered but ties in the data would reduce their validity, and they did not allow modelling of complex interactions. Despite the relatively small number of levels for subject identity, when the standard deviation between subjects could be reliably estimated (excluding near 0 variance), subject identity was implemented as a random factor (Bolker et al., 2009; Gomes, 2022; Oberpriller, de Souza Leite & Pichler, 2022). Considering the nature of the data (strictly positive, right-skewed latencies), a Gamma error distribution (with a log link) was used for modelling.

**Table 3:**
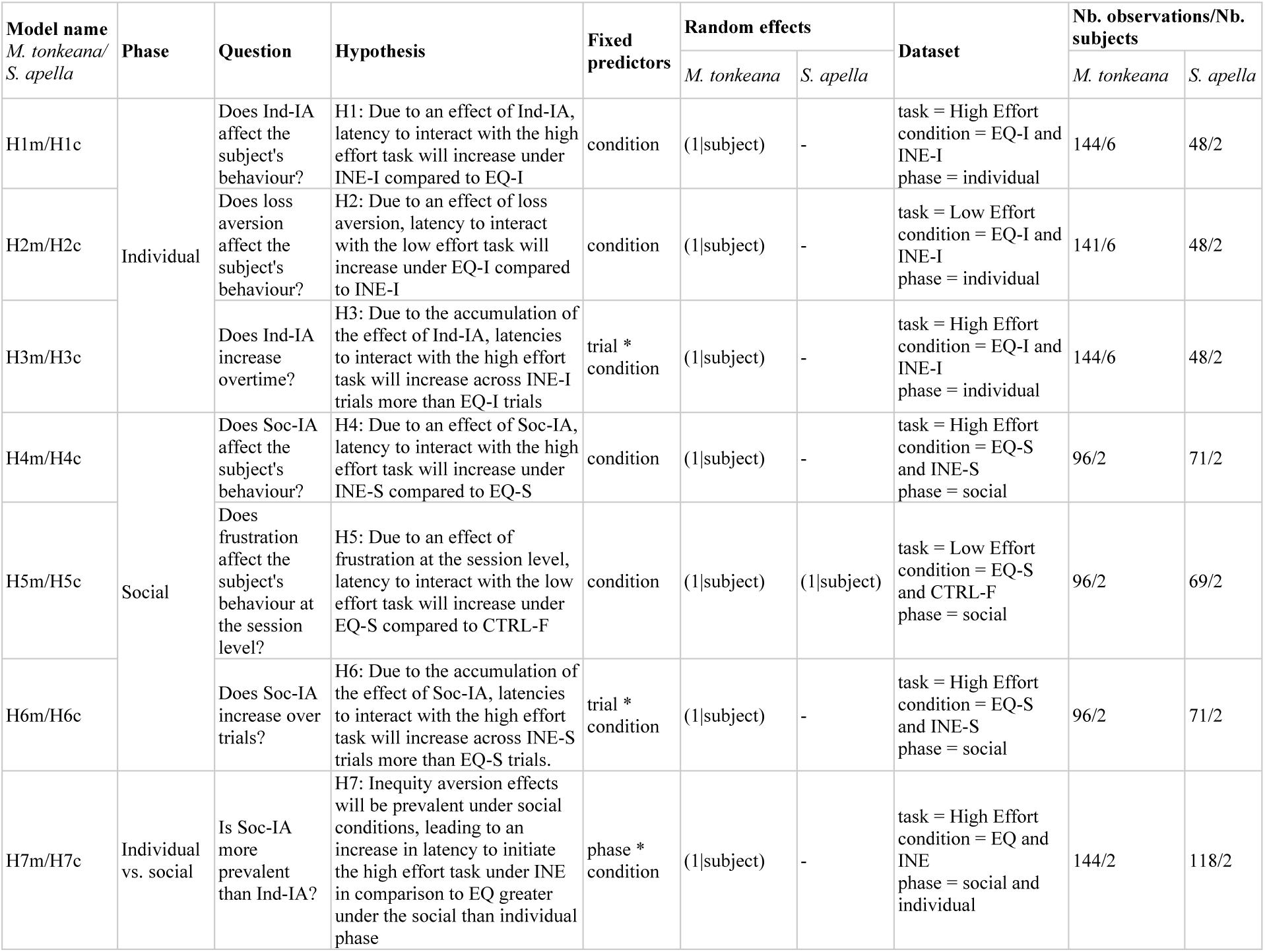
Model specifications for the 7 hypotheses per species. EQ = equity; INE = inequity; I = individual, S = social; CTRL-F = control frustration; Ind-IA = Individually based Inequity aversion; Soc-IA = Socially based Inequity aversion

| Model name<br>*M. tonkeana*/<br>*S. apella* | Phase | Question | Hypothesis | Fixed predictors | Random effects |  | Dataset | Nb. observations/Nb. subjects |  |
| --- | --- | --- | --- | --- | --- | --- | --- | --- | --- |
|  |  |  |  |  | *M. tonkeana* | *S. apella* |  | *M. tonkeana* | *S. apella* |
| H1m/H1c | Individual | Does Ind-IA affect the subject's behaviour? | H1: Due to an effect of Ind-IA, latency to interact with the high effort task will increase under INE-I compared to EQ-I | condition | (1\|subject) | - | task = High Effort condition = EQ-I and INE-I<br>phase = individual | 144/6 | 48/2 |
| H2m/H2c |  | Does loss aversion affect the subject's behaviour? | H2: Due to an effect of loss aversion, latency to interact with the low effort task will increase under EQ-I compared to INE-I | condition | (1\|subject) | - | task = Low Effort condition = EQ-I and INE-I<br>phase = individual | 141/6 | 48/2 |
| H3m/H3c |  | Does Ind-IA increase overtime? | H3: Due to the accumulation of the effect of Ind-IA, latencies to interact with the high effort task will increase across INE-I trials more than EQ-I trials | trial * condition | (1\|subject) | - | task = High Effort condition = EQ-I and INE-I<br>phase = individual | 144/6 | 48/2 |
| H4m/H4c | Social | Does Soc-IA affect the subject's behaviour? | H4: Due to an effect of Soc-IA, latency to interact with the high effort task will increase under INE-S compared to EQ-S | condition | (1\|subject) | - | task = High Effort condition = EQ-S and INE-S<br>phase = social | 96/2 | 71/2 |
| H5m/H5c |  | Does frustration affect the subject's behaviour at the session level? | H5: Due to an effect of frustration at the session level, latency to interact with the low effort task will increase under EQ-S compared to CTRL-F | condition | (1\|subject) | (1\|subject) | task = Low Effort condition = EQ-S and CTRL-F<br>phase = social | 96/2 | 69/2 |
| H6m/H6c |  | Does Soc-IA increase over trials? | H6: Due to the accumulation of the effect of Soc-IA, latencies to interact with the high effort task will increase across INE-S trials more than EQ-S trials. | trial * condition | (1\|subject) | - | task = High Effort condition = EQ-S and INE-S<br>phase = social | 96/2 | 71/2 |
| H7m/H7c | Individual vs. social | Is Soc-IA more prevalent than Ind-IA? | H7: Inequity aversion effects will be prevalent under social conditions, leading to an increase in latency to initiate the high effort task under INE in comparison to EQ greater under the social than individual phase | phase * condition | (1\|subject) | - | task = High Effort condition = EQ and INE<br>phase = social and individual | 144/2 | 118/2 |

Model fit was evaluated using DHARMa simulated residuals (Hartig, 2024) testing for overdispersion, residual uniformity, homoscedasticity, outliers, and zero-inflation. When needed, a separated dispersion model was added to the GLM(M) (Brooks et al., 2017; Nakagawa et al., 2025) (risks of type I error inflation were controlled by verifying that models with heterogeneous variance produced results consistent with models assuming homoscedasticity).When the presence of outliers prevented a good model fit, a sensitivity model without outliers was explored. If outliers accounted for 2% or less of the data, they were removed. When there were multiple outliers and they pertained to a single individual, a model excluding this individual was tested to verify that the significance and direction of the effect remained similar (if so, the complete model was maintained). Specific decisions regarding model construction followed a predefined decision tree (see Supplemental Files Figure S1).

Post-hoc pairwise comparisons of estimated marginal means were conducted using the emmeans package (Lenth, 2025) with Tukey correction for multiple comparisons, with response-scale back-transformation for categorical predictors (condition, phase, preceding task) and estimated slopes (emtrends) for quantitative predictors (trial/session number). Effect sizes for initiation latency were recalculated and summarized as percentages to facilitate comparisons across conditions. Graphical representations were produced using the ggplot2 package (Wickham et al., 2019).

### Data availability statement

All data and code used in this study are available at: https://doi.org/10.5281/zenodo.20626852

## Results

*A summary of all results can be found at Table 4 and the details of all models in Supplemental Files Table S1*.

**Table 4:** Recapitulative view of the effects of the different conditions on the two tasks and species. The direction of the effect (increasing = ↗ or decreasing = ↘) on the latency is indicated by arrows and the magnitude of the effect has been converted to percentages. Results tending to confirm the suspected mechanism at play are in bold.

| Species | Phase | Task | Condition | Effect on latency | Magnitude of effect | Mechanism suspected |
| --- | --- | --- | --- | --- | --- | --- |
| *Macaca tonkeana* | Individual | Low effort | EQ-I compared to INE-I | ↘ | -19% | Loss aversion |
|  |  | High effort | INE-I compared to EQ-I | ↗ | **+49%** | **Individual inequity aversion** |
|  | Social | High effort | INE-S compared to EQ-S | ↗ | **+56%** | **Social inequity aversion** |
|  |  | Low effort | EQ-S compared to CTRL-F | ↗ | **+46%** | **Frustration** |
| *Sapajus apella* | Individual | Low effort | EQ-I compared to INE-I | ↗ | **+44%** | **Loss aversion** |
|  |  | High effort | INE-I compared to EQ-I | ↘ | -83% | Individual inequity aversion |
|  | Social | High effort | INE-S compared to EQ-S | ↗ | **+89%** | **Social inequity aversion** |
|  |  | Low effort | EQ-S compared to CTRL-F | ↗ | **+47%** | **Frustration** |

### 1. Refusals

No refusals to participate in the task or to take or eat the reward was observed. Consequently, this type of response cannot be explored further here.

### 2. Initiation latency

#### 2.1 Individual phase

*H1: Due to an effect of Ind-IA, latency to interact with the high-effort task will increase under INE-I compared to EQ-I*

There was a significant effect of condition on initiation latency of the high-effort task in Tonkean macaques (model H1m: β = 0.41, SE = 0. 17, z = 2.35, p = 0.019) and in capuchins (model H1c: β = -1.79, SE = 0. 42, z = -4.29, p<0.001). Post-hoc pairwise comparisons of estimated marginal means showed that initiation latency was significantly longer under INE-I compared to EQ-I in Tonkean macaques (H1m: EQ-I/INE-I ratio = 0.67, SE = 0.12, z.ratio = -2.35, p = 0.019), whereas capuchins showed significantly shorter initiation latencies under INE-I compared to EQ-I (H1c: EQ-I/INE-I ratio = 5.97, SE = 2.49, z.ratio = 4.29, p < 0.001; Figure 3).

**Figure 3:**
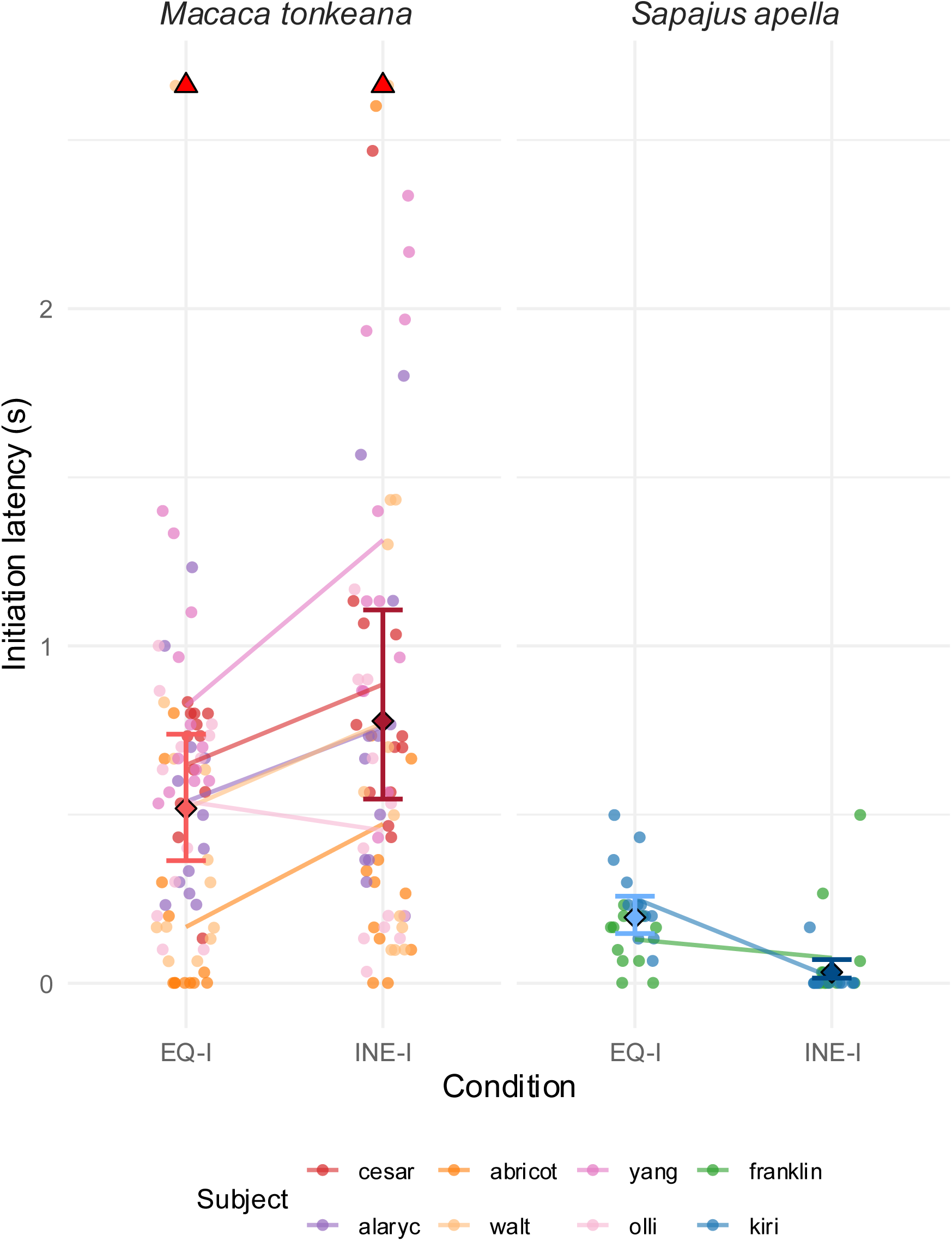
High-effort task initiation latency (in seconds) by condition (EQ-I = individual equity; INE-I = individual inequity) and species in the individual phase. Coloured dots = individual data (n=8); filled diamonds = means ± 95% CI; Red triangles = values above 99th percentile.

*H2: Due to an effect of loss aversion, latency to interact with the low-effort task will increase under EQ-I compared to INE-I*

There was a significant effect of condition on initiation latency of the low-effort task in Tonkean macaques (model H2m: β = 0.21, SE = 0.09, z = 2.22, p = 0.026) and capuchins (model H2c: β = -0.36, SE = 0.13, z = -2.83, p = 0.005). Post-hoc pairwise comparisons of estimated marginal means showed that initiation latency was significantly longer under INE-I compared to EQ-I in Tonkean macaques (EQ-I/INE-I ratio = 0.81, SE = 0.08, z.ratio = -2.22, p = 0.026), whereas capuchins showed significantly shorter initiation latencies under INE-I compared to EQ-I (EQ-I/INE-I ratio = 1.44, SE = 0.19, z.ratio = 2.83, p = 0.005).

*H3: Due to the accumulation of the effect of Ind-IA, latencies to interact with the high-effort task will increase across INE-I trials more than EQ-I trials*

There was no significant effect of trial number on initiation latency of the high-effort task in Tonkean macaques (model H3m: p = 0.998), nor a significant interaction between trial number and condition (p = 0.250). In contrast, in capuchins, there was a significant effect of the interaction between trial number and condition (model H3c: β = 0.52, SE = 0.11, z = 4.79, p < 0.001). Post-hoc pairwise comparisons of estimated marginal means showed that initiation latency significantly increased across trials under INE-I (trial trend = 0.49, SE = 0.10, z.ratio = 4.85, p < 0.001) but not under EQ-I (p = 0.473).

#### 2.2 Social phase

*H4: Due to an effect of Soc-IA, latency to interact with the high-effort task will increase under INE-S compared to EQ-S*

There was a significant effect of condition on initiation latency of the high-effort task in Tonkean macaques (model H4m: β = 0.45, SE = 0.17, z = 2.72, p = 0.007) and in capuchins (model H4c: β = 0.63, SE = 0.18, z = 3.45, p = 0.001). In both species, post-hoc pairwise comparisons of estimated marginal means showed that initiation latency was significantly longer under INE-S compared to EQ-S (H4m: EQ-S/INE-S ratio = 0.64, SE = 0.11, z.ratio = -2.72, p = 0.007; H4c: EQ-S/INE-S ratio = 0.53, SE = 0.97, z.ratio = -3.45, p < 0.001; Figure 4).

**Figure 4:**
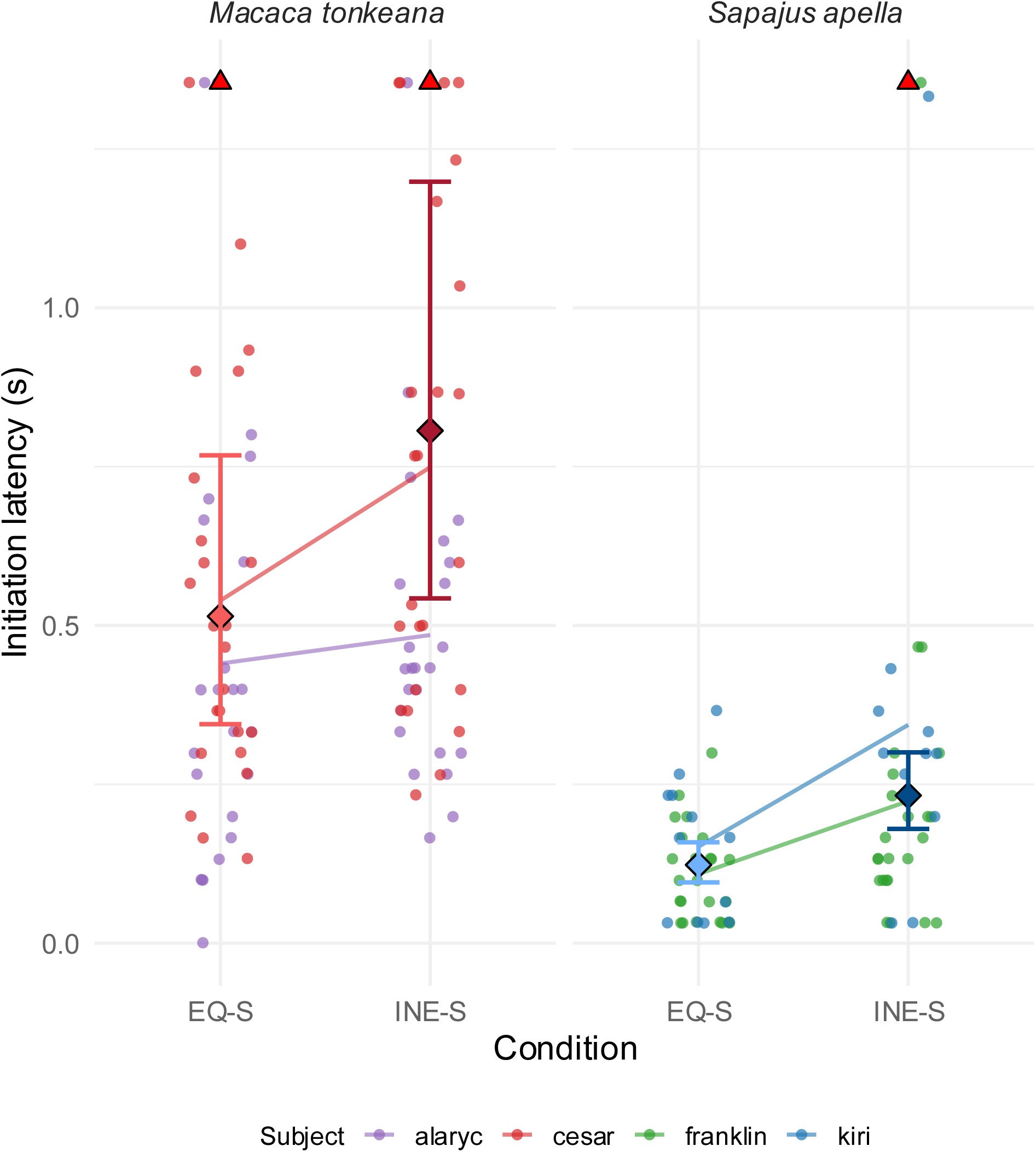
High-effort task initiation latency (in seconds) by condition (EQ-S = social equity; INE-S = social inequity) and species in the social phase. Coloured dots = individual data (n=4); filled diamonds = means ± 95% CI; Red triangles = values above 99th percentile.

*H5: Due to an effect of frustration at the session level, latency to interact with the low-effort task will increase under EQ-S compared to CTRL-F*

There was a significant effect of condition on initiation latency of the low-effort task in Tonkean macaques (model H5m: β = -0.38, SE = 0.08, z = -4.68, p < 0.001) and in capuchins (model H5c: β = -0.38, SE = 0.18, z = -2.18, p = 0.029). In both species, post-hoc pairwise comparisons of estimated marginal means showed that initiation latency was significantly shorter under CTRL-F compared to EQ-S (H5m: EQ-S/CTRL-F ratio = 1.46, SE = 0.12, z.ratio = 4.68, p < 0.001; H5c: EQ-S/CTRL-F ratio = 1.47, SE = 0.26, z.ratio = 2.18, p = 0.029).

*H6: Due to the accumulation of the effect of Soc-IA, latencies to interact with the high-effort task will increase across INE-S trials more than EQ-S trials*.

There was a significant effect of the interaction between trial number and condition on initiation latency of the high-effort task in Tonkean macaques (model H6m: β = 0.13, SE = 0.05, z = 2.75, p = 0.006). Post-hoc pairwise comparisons of estimated marginal means showed that initiation latency became significantly shorter across trials under EQ-S (trial trend = -0.10, SE = 0.03, z.ratio = -3.23, p = 0.002) but not INE-S (p = 0.391). In contrast, in capuchins, there was no significant effect of trial number on initiation latency (p = 0.823), nor a significant interaction between trial number and condition (p = 0.783).

#### 2.3 Social vs. individual phase

*H7: Inequity aversion effects will be prevalent under social conditions, leading to an increase in latency to initiate the high-effort task under INE in comparison to EQ greater under the social than individual phase*

There was no significant effect of phase on initiation latency of the high-effort task in Tonkean macaques (model H7m: p = 0.397), nor a significant interaction between phase and condition (p = 0.586). In contrast, in capuchins, there was a significant effect of the interaction between phase and condition (model H7c: β = 12.61, SE = 0.42, z = 6.27, p < 0.001). Post-hoc pairwise comparisons of estimated marginal means showed that initiation latency was significantly longer under INE during the social phase compared to the individual phase (individual/social ratio = 0.12, SE = 0.034, z.ratio = -7.31, p < 0.001), but phase had no significant effect under EQ (p = 0.132; Figure 5).

**Figure 5:**
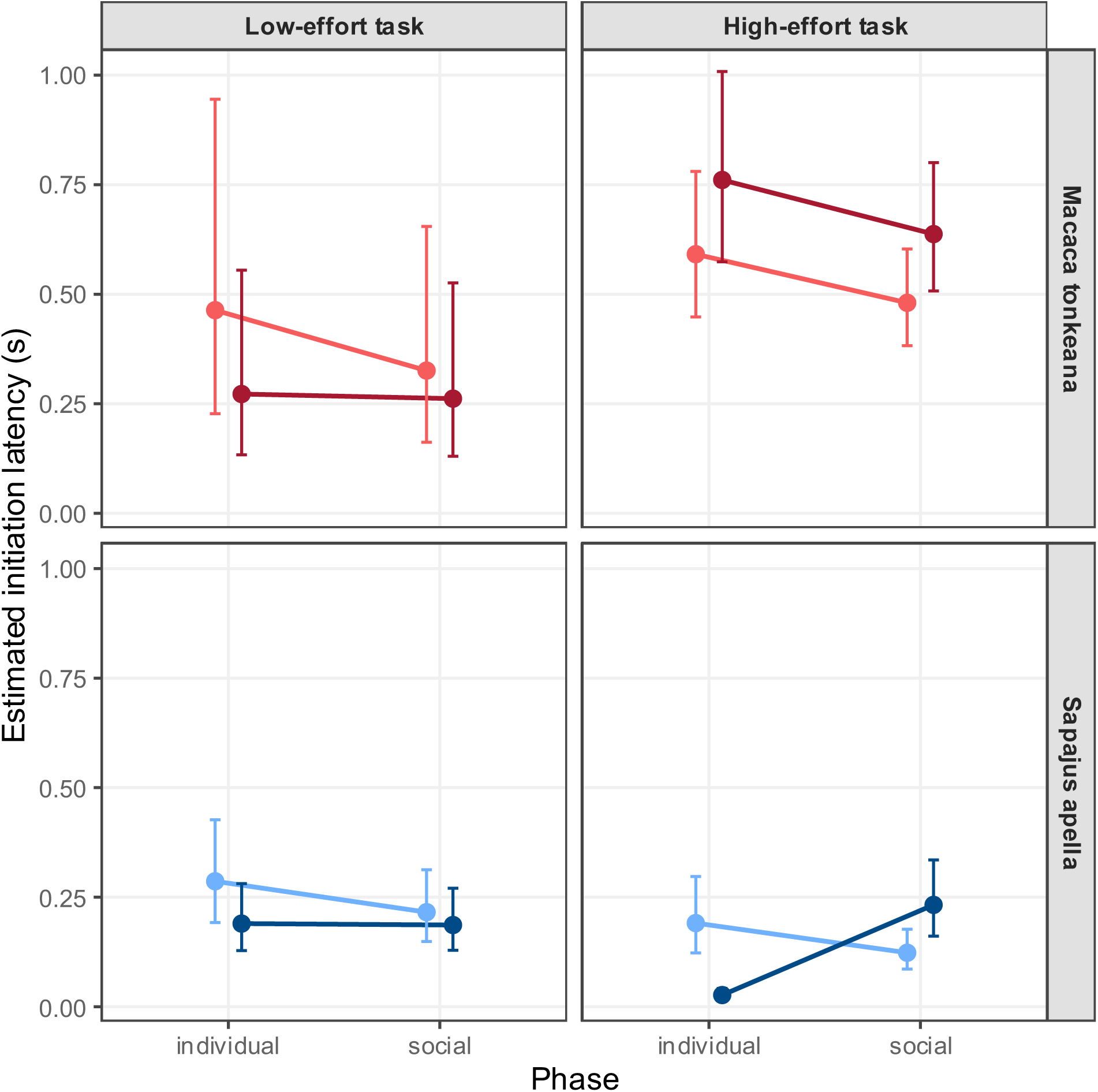
Estimated initiation latencies across phases for Macaca tonkeana and Sapajus apella. Each point shows the estimated mean ± 95% confidence interval for the individual and social phases. Lighter colours indicate EQ conditions and darker colours indicate INE conditions. Specifically, light blue = capuchin EQ, dark blue = capuchin INE, light red = macaque EQ, dark red = macaque INE. Facets separate species (rows) and task type (columns: high-effort vs. low-effort).

## Discussion

This study aimed to evaluate the potential of a new methodological approach to demonstrate the existence of socially (Soc-IA) and individually (Ind-IA) driven *inequity aversion*, while controlling for *individual contrast* effects (Ind-C). To do so, we presented subjects with two tasks requiring different levels of effort; which they had learned would result in two specific reward quantities. We then manipulated the rewards so that individuals sometimes received less than they deserved relative to their effort (creating potential Ind-IA) or less than their partner (creating potential Soc-IA).

Throughout this study, individuals showed no refusals to perform the task or accept the reward, regardless of the condition. However, inequity significantly affected their latency to engage with the task. During the individual phase, both species exhibited distinct patterns. Macaques took longer to engage with the high-effort task under conditions of inequity while capuchins were faster. Capuchins also showed an increase in latency for the low-effort task under conditions of equity, consistent with a loss aversion effect. In the social phase, inequity elicited a convergent response in both species, characterised by a slower engagement with the high-effort task. Taken together, these patterns suggest that sensitivity to inequity may rely on both social and non-social processes in macaques, whereas it appears more closely linked to the social context in capuchins. With those results in mind, the present study remains primarily exploratory in nature. Given the small sample size, its main objective is to assess the feasibility of a new methodological approach. Consequently, the following discussion focuses mainly on how alternative hypotheses were addressed, the limitations of these controls, and how the protocol could be improved in the future.

### Methodological approach: latencies and quantities

The time taken to engage in the task is a more subtle and less overtly performative behaviour than refusal. However, it has been shown that latencies are a good indicator of emotional state and stimulus impact (Watanabe et al., 2001; Gordon & Rogers, 2015; Bethell et al., 2016; Blanchette et al., 2017), making them particularly interesting and valid measures for testing *inequity aversion*. Since no refusals were recorded in this study, the interpretation of the effect of inequity relied entirely on the latency to engage with the task. This distinguishes our study from many others on *inequity aversion* (e.g. van Wolkenten, Brosnan & de Waal, 2007; Talbot et al., 2011; Sosnowski et al., 2021), although some research has already incorporated these measures (Brosnan & de Waal, 2003; Silberberg et al., 2009; Titchener et al., 2023).

The absence of refusals observed here could be explained by several factors. Firstly, previous studies have shown that using a difference in food quantity does not always elicit refusal where a difference in quality would (Feller, 2016; Talbot et al., 2018 but see: Fletcher, 2008; Feller, 2016). This was a potential limitation considered during the design of our methodology, but several advantages associated with using quantity rather than quality motivated our choice. Indeed, differences in the quality of rewards, such as a grape *versus* a cucumber, may not hold the same subjective value for all individuals. This remains true even when all express a general preference for one over the other. In contrast, differences in the quantity of a valued reward are more likely to be perceived similarly by all subjects. Using quantity rather than quality also allowed us to conduct tests over several weeks and successive trials while maintaining a stable comparative value between rewards. This is more difficult to achieve when using two rewards of different qualities (Addessi, 2008; Martin et al., 2018; Vonk, Truax & McGuire, 2022). From an ecological perspective, sensitivity to quantitative inequity alone also seems particularly relevant: in the wild, the distribution of resources often involves the sharing of a single type of resource, such as portions of meat after a hunt (Boesch, 1994).

Another explanation for the lack of refusals is that our methodology involved choices of relatively low stakes for the individuals. This was necessary to ensure sustained participation, as increasing the difficulty of the task would have risked causing the individuals to lose interest over time. It also helped avoid ceiling effects, as a reward that was too high might never have been refused or perceived negatively (Talbot et al., 2018).

### Individually based Inequity Aversion

The slower response time exhibited by macaques when offered an inequitable reward during the individual phase cannot be fully explained by a simple loss aversion mechanism. Indeed, when tested for their reaction to loss aversion as non-task-specific, they did not show the same trend as the one observed in inequity situations. Since frustration was not elicited differently across conditions during the individual phase due to its design, frustration alone cannot explain the reaction either. These results are consistent with the idea that the macaques tested in this study relied on internally defined effort-reward expectations in the absence of a social comparison. It is also worth noting that the effect of inequity did not vary across trials, suggesting a stable response rather than learning or fatigue effects, contrary to what had previously been observed in long-tailed macaques who exhibited longer latencies across trials and sessions (Titchener et al., 2023). In contrast, the capuchins’ faster engagement under individual inequity suggests that factors other than individually defined standards of equity may have been the primary drivers of subjects’ behaviour. When tested for general loss aversion effects, capuchins showed slower initiation latencies for the low-effort task when a higher-value reward was distributed in the session. This sensitivity to loss may have impacted the entire session, which would also explain the slower engagement observed for the equitably rewarded high-effort task as well. These results tend to confirm the predominance of loss aversion in these subjects’ responses to an inequitable payoff. This is consistent with some previous research highlighting capuchins’ sensitivity to contrast effects (Roma et al., 2006; Chen, Lakshminarayanan & Santos, 2006; Rocha et al., 2020, but see Silberberg et al., 2008). That said, these results do not completely rule out the existence of an individually based inequity averse mechanism. Given the increase in latency across inequitable trials, it is possible that inequity had a cumulative effect here. Such an effect has already been observed in capuchins: Brosnan and de Waal (2003) found an increase in refusals to exchange during the latest trials of a session. It is interesting to note that, upon examining latencies, they also observed that capuchins made exchanges more quickly under inequitable condition, even though the overall number of refusals was higher. This suggests that faster engagement does not necessarily reflect acceptance and may, in some cases, be compatible with a negative assessment of the outcome.

### Socially based Inequity Aversion

During the social phase, capuchins did show longer latencies to engage with the inequitably rewarded tasks, which was not the case in the individual phase. While it is possible that reactions to discomfort vary depending on the presence or absence of a partner, these subtle changes in latency likely reflect decisions driven by internal affective states (Gordon & Rogers, 2015; Bethell et al., 2016; Blanchette et al., 2017) rather than performative decisions. If aversion to inequity operates similarly at the individual and social levels, our results suggest that capuchins may use different mechanisms depending on the social context. During the social phase, capuchins also initiated the low-effort task more slowly in equity conditions compared to sessions where no distribution of a higher reward took place. However, this effect was only about half the magnitude of that observed for tasks rewarded inequitably. Given this, as well as the absence of loss aversion effects by design during this phase, the longer response time observed in situations of social inequity were likely not solely due to individual contrast effects. These findings are therefore consistent with the hypothesis that the capuchins reacted primarily to the inequity of the situation, using their social partner as their main reference.

Regarding the macaques, a slower engagement under social inequity was also observed compared to situations of equity. This effect did not differ significantly from that induced by individual inequity. This suggests that, overall, the macaques’ performance was not influenced by the presence of a partner, which does not support the conclusion that social comparison reinforces *inequity aversion*. Moreover, although trial-related effects were observed under equity conditions, this was not the case for social inequity. This stable response pattern was similar to that observed under individual conditions. Similarly to capuchins, macaques displayed frustration, as they were faster to engage with the low-effort task if no equitably rewarded high-effort task was available. That said, this effect was also slightly smaller in magnitude than the effect of social inequity. The control condition used to assess frustration also involved only a low level of effort, which may have made the session less cognitively and physically demanding overall, thereby facilitating faster reaction times. Given this trend, the observed frustration effect could be consistent with the presence of socially based *inequity aversion* in macaques.

While capuchin monkeys showed very different responses to the individual and social phases, Tonkean macaques’ behaviour remained constant. This interspecific contrast might be explained by capuchins’ usually higher levels of activity and reactivity (Fragaszy, Visalberghi & Fedigan, 2004) and their greater impulsivity in cognitive tasks (Pelé et al., 2011). They may therefore favour more readily accessible mechanisms, such as loss aversion, in non-social contexts. In contrast, Tonkean macaques’ tolerance for delayed engagement and sustained attention (Pelé et al., 2010; Loyant et al., 2023) might give them the time and resources necessary to mobilize expectations based on personal effort. It is also possible that capuchins require more iterations of inequity to elicit an aversion, as they did show increased latency to engage by the end of individual inequitable sessions. An alternative hypothesis is that the subjective assessment of effort may have differed between the capuchins and macaques tested here. For the capuchins, the energetic or cognitive cost associated with the high-effort task may have been lower (Visalberghi, 2008), thereby reducing the influence of personal merit as a reference point.

### Some limitations

#### Association between task and reward

To confirm the results obtained during the individual phase, it is important to validate that the subjects had stable expectations regarding the reward for each task. However, we did not have a measure that directly tested this association. That said, the data showed that latencies for the low-effort task were either shorter or showed no significant difference when it was the only task in the session, compared to when it was alternated with the high-effort task (rewarded with four raisins). This pattern suggests that subjects did not lose their motivation when they repeatedly received low-value rewards, indicating that they maintained a constant expectation of payoff for the low-effort task. If subjects had perceived the rewards as random and had not understood the relationship between the task and the reward, one might have expected a decline in motivation when only low rewards were available, driven by the expectation of receiving a better payoff during the session (Papini et al., 2022). This hypothesis is further supported in macaques by their selective response observed during the high-effort task. Subjects took longer to engage with the high-effort task when given a single raisin under individual inequitable conditions and were faster in the low-effort task under individual equitable conditions. If rewards were perceived as random, one would expect a uniform negative reaction to a lower reward, reflecting a general aversion to loss, for both tasks and sessions (Oberliessen & Kalenscher, 2019; Ritov et al., 2024). But since the longer latencies were selectively associated with inequity, this suggests that the macaques expected a specific reward depending on the task (Ott, Stein & Nieder, 2023), which supports the idea that the association between the task and the reward was well understood.

#### Motivation

One limitation of the individual phase of this study was the inability to establish a motivational baseline for the high-effort task when the reward was limited to a single raisin. During the motivational control, we believed that subjects had already learned to perform the task efficiently. Unfortunately, this was not the case, as their performance later improved considerably. As a result, the data from this initial phase were no longer comparable to those collected during subsequent test sessions and could not serve as a valid motivational baseline. As for the capuchins, since the results showed faster engagement under inequitable condition, this does not constitute a major concern. Regarding the macaques, they exhibited longer engagement latencies for the low-effort task under inequitable conditions. This suggests that the inequitable payoff offered for the high-effort task had a carryover effect (Papini, 2014) that impacted the low-effort task (which was rewarded equitably). If the change in latencies reflected a simple targeted lack of motivation, there is no reason to believe it would have led to a negative-valance emotion affecting the low-effort task, as aversion to inequity might. That said, while these factors provide indirect evidence that the motivation effect may be excluded, they do not allow us to rule it out at this stage.

#### Expectation violation

Another alternative explanation for our positive findings on individually based *inequity aversion* is the potential violation of expectations regarding the presentation of the task. After seeing a single reward, subjects would initially expect to be offered the task requiring low effort. When presented with the high-effort task, longer latencies might reflect the time needed to adapt their behaviour to an unexpected task rather than a reaction to inequity per se (Tinklepaugh, 1928). Although this possibility cannot be entirely ruled out, the methodology was designed to limit such effects. After the reward was displayed, the task apparatus (the wooden ball) was briefly presented next to the reward before the task began. This was intended to allow for behavioural adjustment and to reduce any surprise effect by enabling subjects to anticipate the upcoming task before engaging in it.

#### The sample

Another point that calls for great caution in this study concerns the sample size. The protocol required extensive training and specific setups for social testing. It also prevented the monkeys that served as models and received the highest reward during the social phase from subsequently being tested as subjects. These constraints limited the number of individuals that could be comprehensively tested and included in the analysis. As a result, all interpretations made in this study must be limited to the tested subjects and cannot be generalized to the species or population level. Moreover, due to the limited number of individuals, personal characteristics such as social rank, sex, age, or temperament, which are known to influence reactions to inequity (Brosnan & de Waal, 2003; Brosnan et al., 2010, 2015; Amici, Call & Aureli, 2012; Hopper et al., 2013; Mustoe et al., 2016; Yasue et al., 2018), could not be explored. Only adult males were tested, which ensured sample consistency but prevented analysis of these additional factors. Based on these results, we primarily aim to discuss the potential of this methodological shift and the consideration of individual merit, while highlighting limitations and future directions.

#### Future directions

Future research should aim to use measures that can be directly compared across tasks to facilitate the testing of alternative hypotheses. This could be achieved by developing latency measures that can be standardized across task types for example.

Future studies would also benefit from implementing a formal assessment of the animals’ understanding of the task-reward relationship before and after the tests. Such controls would help dispel certain doubts regarding the mechanisms underlying the animals’ response, ensuring that this understanding is maintained throughout the sessions. Similarly, it is essential to establish, prior to introducing inequity, a reliable baseline for motivation regarding the performance of the high-effort task in exchange for a lower value reward. In the present study, this control failed due to the longer-than-expected time it took for the subjects to reach a consistent efficiency. We therefore recommend extending training by several sessions, even when subjects reach an apparent plateau, before getting to the next stage.

Future research aiming at incorporating individual merit into studies on *inequity aversion* should also prioritise protocols that control for social status and, ideally, other individual characteristics such as sex, personality or age. Given the established influence of hierarchy on reactions to food distribution (Amici, Call & Aureli, 2012), failing to control for this variable could skew the results. Exploring this paradigm with tolerant primate species also appears promising, as previously theorized (Brosnan, 2006), since *inequity aversion* might be more pronounced in these species. As these exploratory findings suggest, it might even exist in an individual context. Repeating and extending this experiment with capuchins would also be informative, as an effect of the repetition of inequity was observed during the individual phase. It is therefore possible that individually based inequity sensitivity exists in capuchins but that it only manifests after a greater number of repetitions.

## Conclusions

This study aimed to determine whether the use of a new methodology, involving different tasks and rewards, could reveal the existence of an individually based *inequity aversion*. In both species tested, latencies to engage with the tasks were reliably altered according to the different test conditions. In the individual phase, macaques took longer to engage with the inequitably rewarded task, a behaviour that cannot be explained by *individual contrasts*. Capuchins, on the other hand, showed a behaviour more consistent with loss aversion effects. Under social conditions, both species displayed slower engagement when the outcome was inequitable. It is important to note that these analyses remain exploratory and are limited at this stage to the tested individuals. The small sample size precludes generalization and the consideration of sociodemographic factors. The lack of a motivational baseline also warrants caution in the interpretation of the results. Finally, although several indications suggest that subjects understood the association between the task and the reward, this was not tested directly and cannot be definitively established within the scope of the present study. Despite these limitations, this study provides initial evidence that manipulating effort and reward within a task could be a promising methodological approach for studying *inequity aversion* beyond a strict social framework. Future studies, drawing on larger and more diverse samples, incorporating explicit tests of understanding of the link between task and reward, a measure of engagement comparable across tasks and robust motivational baselines will be necessary to determine whether non-social mechanisms of *inequity aversion* are truly at work in primates.

## Acknowledgments

The authors are grateful to the Simian Laboratory Europe (Silabe, University of Strasbourg, France) for supporting this research and welcoming their team. They warmly thank the animal care and welfare staff for their valuable assistance throughout the project. They also sincerely thank the interns for their commitment and important contributions to this study: Ninon Chautard, Renée Massip, Zazie Benoit-Delaby, and Sarah Piaugeard for their help with the experimental procedures, with the latter two also contributing to video coding. A special thank you to Manon Serda for the realisation of the illustrations present in this article.

