## Supplemental Information S1 for "Social or non-social? An exploratory approach to study inequity aversion in primates"

***Supplemental Information S1: Control for arousal effects***

### Control for arousal effects:

In the social inequity condition, the presence of a large amount of food (eight raisins) given to the partner could potentially trigger heightened arousal in the subject. This could lead to altered behaviour, regardless of any perceived inequity.

To control for this, we conducted two additional sessions per tested pair after all the equity and inequity sessions had been completed. In this control (CTRL-Q), the subject received eight raisins for the high-effort task in a setup identical to that used in the social equity condition. This allowed us to control for the reaction to a large amount of food presented in an equitable context and to assess whether the increased arousal due solely to the presentation of the reward could explain the observed behavioural patterns.

### Results:

*Hypothesis: Due to an arousal effect, latency to interact with the high-effort task will increase under CTRL-Q conditions in comparison to EQ-S conditions*

There was no significant effect of condition on initiation latency of the high-effort task in Tonkean macaques (model H6m: $p = 0.065$). In contrast, in capuchins, there was a significant effect of condition on initiation latency (model H6c: $\beta = 0.49$, $SE = 0.17$, $z = 2.86$, $p = 0.004$). Post-hoc pairwise comparisons of estimated marginal means showed that initiation latency was significantly longer under CTRL-Q compared to EQ-S in capuchins (EQ-S/CTRL-Q ratio = 0.61, $SE = 0.11$, z.ratio = -2.86, $p = 0.004$), corresponding to an increase of 64%.

### Discussion:

During the social inequity phase, a large number of rewards were clearly visible and distributed. After controlling for the effect of arousal, results showed that this did not alter macaques' behaviour. But for capuchins, even in the absence of inequity, the use of such quantities led to a slowing of responses. Arousal or distraction may therefore have occurred, delaying engagement in the task. However, its magnitude was smaller than that of the social inequity effect. Results therefore remain consistent with a socially based inequity hypothesis for the capuchins as well as for the macaques.
