## Supplementary figures and images for "Social or non-social? An exploratory approach to study inequity aversion in primates"

### Supplemental Figure S1

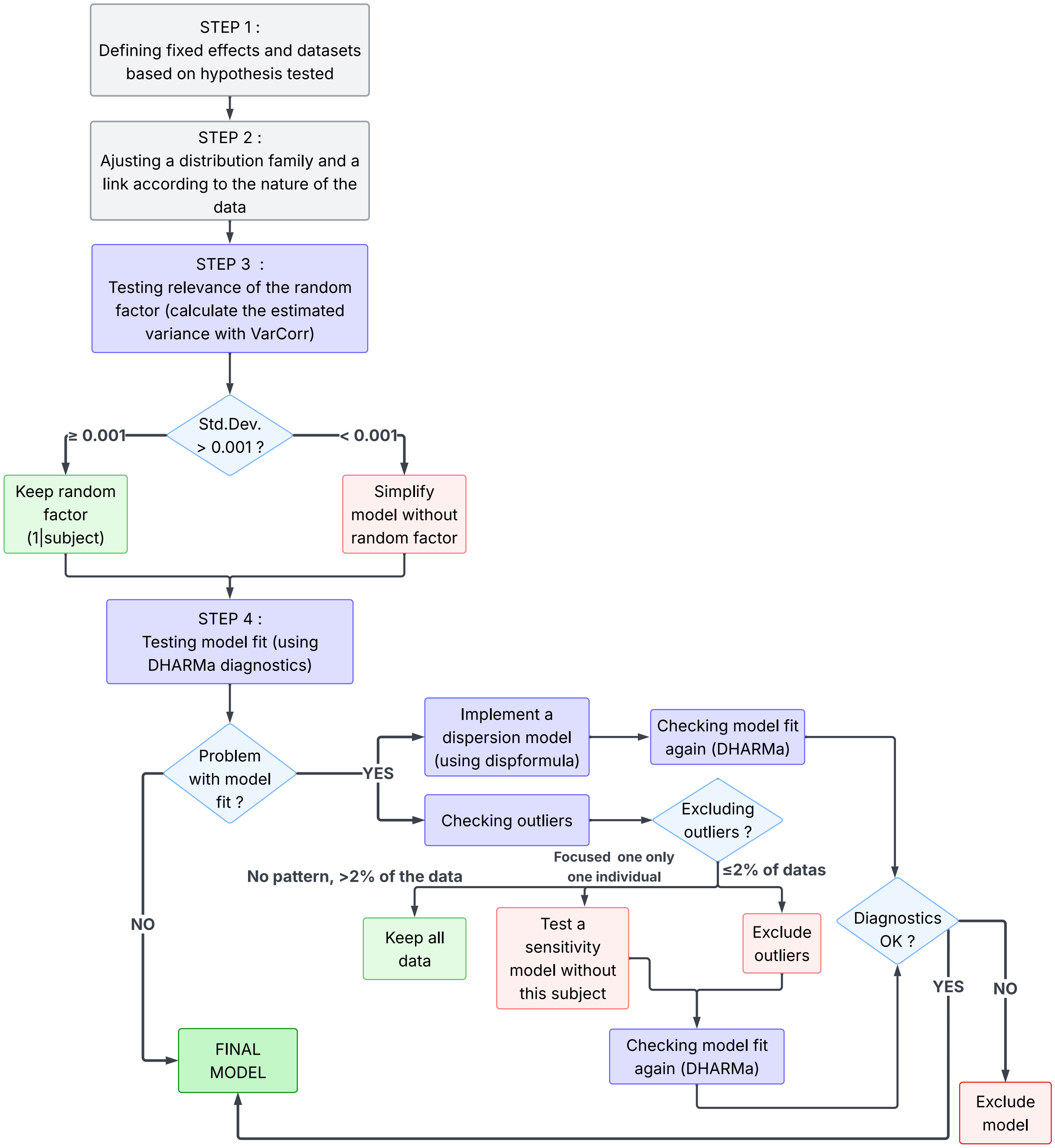


***Supplemental Figure S1: Decision tree for statistical model building***
