## Supplemental Table S1 for "Social or non-social? An exploratory approach to study inequity aversion in primates"

***Supplemental Table S1: Summary of the results of all statistical models***.
*Parameter estimates, standard error, z value, and p-values are shown for fixed effects.*

| Predictors | Estimates | std. Error | z value | p |
|---|---|---|---|---|
| H1m: _initiation ~ condition + (1 \| subject)_ | | | | |
| condition [INE] | 0.41 | 0.17 | 2.34 | **0.019** |
| H1c: _initiation ~ condition_ | | | | |
| condition [INE] | -1.79 | 0.42 | -4.29 | **<0.001** |
| H2m: _initiation ~ condition + (1 \| subject)_ | | | | |
| condition [INE] | 0.21 | 0.09 | 2.22 | **0.026** |
| H2c: _initiation ~ condition_ | | | | |
| condition [INE] | -0.36 | 0.13 | -2.83 | **0.005** |
| H3m: _initiation ~ trial * condition + (1 \| subject)_ | | | | |
| trial | 0.00 | 0.04 | 0.00 | 0.998 |
| condition [INE] | 0.78 | 0.38 | 2.05 | **0.040** |
| trial * condition [INE] | -0.06 | 0.05 | -1.15 | 0.250 |
| H3c: _initiation ~ trial * condition_ | | | | |
| trial | -0.03 | 0.04 | -0.72 | 0.473 |
| condition [INE] | -5.84 | 0.74 | -7.88 | **<0.001** |
| trial × condition [INE] | 0.52 | 0.11 | 4.79 | **<0.001** |
| H4m: _initiation ~ condition + (1 \| subject)_ | | | | |
| condition [INE] | 0.45 | 0.17 | 2.72 | **0.006** |
| H4c: _initiation ~ condition_ | | | | |
| condition [INE] | 0.63 | 0.18 | 3.45 | **0.001** |
| H5m: _initiation ~ condition + (1 \| subject)_ | | | | |
| condition [CTRL-F] | -0.38 | 0.08 | -4.68 | **<0.001** |
| H5c: _initiation ~ condition + (1 \| subject)_ | | | | |
| condition [CTRL-F] | -0.38 | 0.18 | -2.18 | **0.029** |
| H6m: _initiation ~ condition_ | | | | |
| condition [CTRL-Q] | 0.07 | 0.16 | 0.46 | 0.645 |
| H6c: _initiation ~ condition_ | | | | |
| condition [CTRL-Q] | 0.49 | 0.17 | 2.86 | **0.004** |
| H7m _initiation ~ trial * condition + (1 \| subject)_ | | | | |
| trial | -0.10 | 0.03 | -3.12 | **0.002** |
| condition [INE] | -0.44 | 0.38 | -1.18 | 0.239 |
| trial * condition [INE] | 0.13 | 0.05 | 2.75 | **0.006** |
| H7c: _initiation ~ trial * condition_ | | | | |
| trial | -0.01 | 0.04 | -0.22 | 0.823 |
| condition [INE] | 0.52 | 0.47 | 1.11 | 0.268 |
| trial * condition [INE] | 0.02 | 0.06 | 0.28 | 0.783 |
| H8m: _initiation ~ phase * condition + (1 \| subject)_ | | | | |
| phase [social] | -0.15 | 0.18 | -0.85 | 0.397 |
| condition [INE] | 0.33 | 0.21 | 1.60 | 0.109 |
| phase [social] * condition [INE] | 0.14 | 0.25 | 0.55 | 0.586 |
| H8c: _initiation ~ phase * condition_ | | | | |
| phase [social] | -0.44 | 0.29 | -1.50 | 0.132 |
| condition [INE] | -1.97 | 0.32 | -6.12 | **<0.001** |
| phase [social] * condition [INE] | 2.61 | 0.42 | 6.27 | **<0.001** |
